# Neurofeedback Training on Motor Cancellation Enhances Peripheral but not Cortical Beta Band Oscillations

**DOI:** 10.64898/2026.08.11.743916

**Authors:** Emanuele Abbagnano, Benedetta Memè, Alejandro Pascual-Valdunciel, Yongkun Zhao, Jaime Ibáñez, Dario Farina

## Abstract

Beta oscillations (13–30 Hz) are a prominent sensorimotor neural rhythm and an important biomarker in neurorehabilitation. These oscillations propagate along the corticospinal pathway and are expressed in the discharge patterns of spinal motor neurons, enabling the assessment of corticomuscular coupling. Moreover, peripheral beta band oscillations have recently emerged as a potential control signal for motor augmentation interfaces. However, it remains unclear whether peripheral beta activity simply reflects cortical oscillations or is partly shaped by peripheral mechanisms, and to what extent it can be voluntarily controlled. To address these questions, we developed a 10-day neurofeedback protocol in which participants learned to up-regulate peripheral beta band activity. Subjects were trained to exploit movement cancellation, a behaviour naturally associated with increased cortical and muscle beta band activity, as a two-state strategy to voluntarily modulate peripheral beta band power. Each session included a guided familiarization phase based on a GO/NO-GO task, in which participants familiarized with movement cancellation through guided visual cues, followed by an asynchronous control phase in which they self-initiated the same strategy without external guidance to increase peripheral beta band activity in a target window. Participants progressively improved their ability to voluntarily modulate peripheral beta band activity. Peripheral beta band power during movement cancellation increased significantly across training days in both the familiarization and asynchronous control phases. Intramuscular coherence in the beta band also increased, indicating enhanced common synaptic input to the motor neuron pool in this band. In contrast, cortical beta power and corticomuscular coherence remained unchanged. Together, these findings demonstrate that peripheral beta band activity is a dynamic neural feature that can be voluntarily shaped through training, supporting its potential as a non-invasive control signal for future neurorehabilitation and motor augmentation technologies.

## Introduction

Beta band neural oscillations (13-30 Hz) are linked to sensorimotor control within the sensorimotor cortex (Baker, 2007; Kilavik et al., 2013). Alterations in beta band activity are closely linked to motor dysfunction in neurological conditions such as Parkinson’s disease, stroke, and spinal cord injury (Gourab and Schmit, 2010; Von Carlowitz-Ghori et al., 2014; Zokaei et al., 2021). Moreover, beta band activity has been successfully used as a control signal for closed-loop deep brain stimulation, where stimulation is delivered in response to elevations in beta power to suppress pathological neural synchronization in Parkinson’s disease (Tinkhauser et al., 2017). Together, these findings highlight the value of beta band activity as both a biomarker and therapeutic target in neurorehabilitation, underscoring the need for accessible and non-invasive methods to monitor these neural signals.

These oscillations are transmitted uniformly across the motor neuron pool, independently of motor neuron size, and reach the muscles through the fastest corticospinal pathways (Abbagnano et al., 2025; Ibáñez et al., 2021). Because common synaptic input is linearly transmitted from the motor neuron pool to the muscle (Farina and Negro, 2015), beta band activity can be decoded from motor unit discharge patterns using muscle recordings (Ibáñez et al., 2025). During sustained motor tasks such as isometric contractions, cortical and peripheral beta band activity exhibit strong functional coupling, suggesting a predominantly cortical origin of the peripheral signal (Baker, 2007; Conway et al., 1995; Echeverria-Altuna et al., 2022; Witham et al., 2011). Consistent with this relationship, real-time neurofeedback of peripheral beta band activity enables voluntary modulation of beta band amplitude, accompanied by corresponding changes in cortical beta activity (Bräcklein et al., 2022). This makes peripheral beta band activity a promising target for non-invasive neural interfaces capable of accessing brain-derived beta oscillations from muscle recordings (Ibáñez et al., 2025). These interfaces could provide a practical and accessible approach for applications in both neurorehabilitation and motor augmentation.

However, several challenges remain for the development of such interfaces. Evidence from studies of spinal inhibitory circuitry suggests that beta band activity recorded in tonically active muscles is not merely a reflection of cortical beta oscillations, but is also shaped by spinal processing (Dernoncourt et al., 2025; Matsuya et al., 2017; Williams and Baker, 2009a). In particular, recurrent inhibitory circuits mediated by Renshaw cells can interact with common synaptic input in the beta band and influence the frequency content of motor neuron output, thereby modulating synchronized activity at the motor unit level (Williams and Baker, 2009b). Consequently, peripheral beta band activity may reflect not only descending cortical drive but also its interaction with spinal mechanisms, in particular in tasks characterized by inhibitory signals.

To further investigate the relation between cortical and peripheral beta oscillations, we developed a 10-day longitudinal neurofeedback protocol designed to improve voluntary control of peripheral beta band activity by motor cancellation. Participants were trained to exploit the increase in peripheral beta band activity associated with movement cancellation as a two-state control strategy, while concurrent EEG and HD-EMG recordings were acquired before and after training and used to determine the site of adaptation (peripheral/cortical). We hypothesized that improved capacity to cancel a task would be accompanied by increased peripheral beta band activity that could be explained either with changes in spinal mechanisms or with a concurrent increase in cortical beta. We found that the motor cancellation training enhanced peripheral beta band activity and common synaptic input to the motor neuron pool without detectable changes in cortical beta band activity or corticomuscular coherence, suggesting that the adaptations arose predominantly from peripheral and spinal mechanisms.

## Methods

### Experimental Data Acquisition

#### Subjects

Eight healthy participants (ages: 24.87 ± 1.88 years, 4 males and 4 females) were recruited for this study. All participants provided written informed consent prior to their inclusion. The study was conducted in accordance with the principles outlined in the Declaration of Helsinki and received approval from the Imperial College London Ethics Committee (reference number: 18IC4685).

#### Data Acquisition

High-density surface EMG was recorded from the tibialis anterior muscle of the dominant leg only before and after the 10-day longitudinal training period. Signals were acquired using a 256-channel grid (26 rows × 10 columns; gold-coated electrodes, 1 mm diameter, 4 mm inter-electrode distance; OT Bioelettronica). The grid was centered over the muscle belly and oriented along the fiber direction. Signals were recorded in a monopolar configuration and amplified with the Quattrocento system (OT Bioelettronica, Torino, Italy). Data were sampled at 2048 Hz and processed with a digital band-pass filter from 10 to 500 Hz.

Bipolar EMG signals were recorded from the tibialis anterior muscle of the dominante leg throughout the 10-day longitudinal training period using four disposable surface electrodes (Ambu Neuroline 720) arranged as two longitudinal pairs. The first pair was placed over the muscle belly with a few mm of inter-electrode spacing, and the second pair was positioned 2 cm distal to it. A reference electrode was placed around the ankle using a wet band in contact with the bone. Signals were acquired with a Quattrocento amplifier at 2048 Hz using a bipolar cable (AD8x2JD, OT Bioelettronica).

Ankle dorsiflexion force was measured via a load cell (TF-022, CCT Transducer s.a.s) affixed to the dynamometer pedal and digitized with the Quattrocento Amplifier system at 2048 Hz.

Electroencephalography (EEG) was collected using 31 active gel electrodes arranged according to the 10–20 layout (actiCAP, Brain Products GmbH), with FCz selected as the reference during acquisition. Signals were amplified with the BrainVision actiCHamp Plus amplifier, initially sampled at 1000 Hz, and later resampled to 2048 Hz for alignment with the EMG data. Synchronization across all devices was achieved using a common digital trigger delivered to both the Quattrocento and actiCHamp Plus systems.

### Experimental Design

We examined whether participants could improve voluntary control of peripheral beta band activity, and whether the resulting behavioural adaptations were associated solely with changes in peripheral beta band activity or also with changes in cortical beta band activity.

The protocol consisted of two parts (Figure 1A). First, participants completed a 10-day longitudinal training period focused on active familiarization and voluntary modulation of peripheral beta activity using bipolar EMG as neurofeedback. Second, we assessed potential training-related changes in the corticospinal transmission of beta oscillations using concurrent EEG and high-density surface electromyography (HD-EMG) recordings collected before and after the training. During these sessions, participants performed isometric contractions, a condition known to reliably enhance beta band power at both cortical and peripheral levels (Baker, 2007; Engel and Fries, 2010).

**Figure 1.**
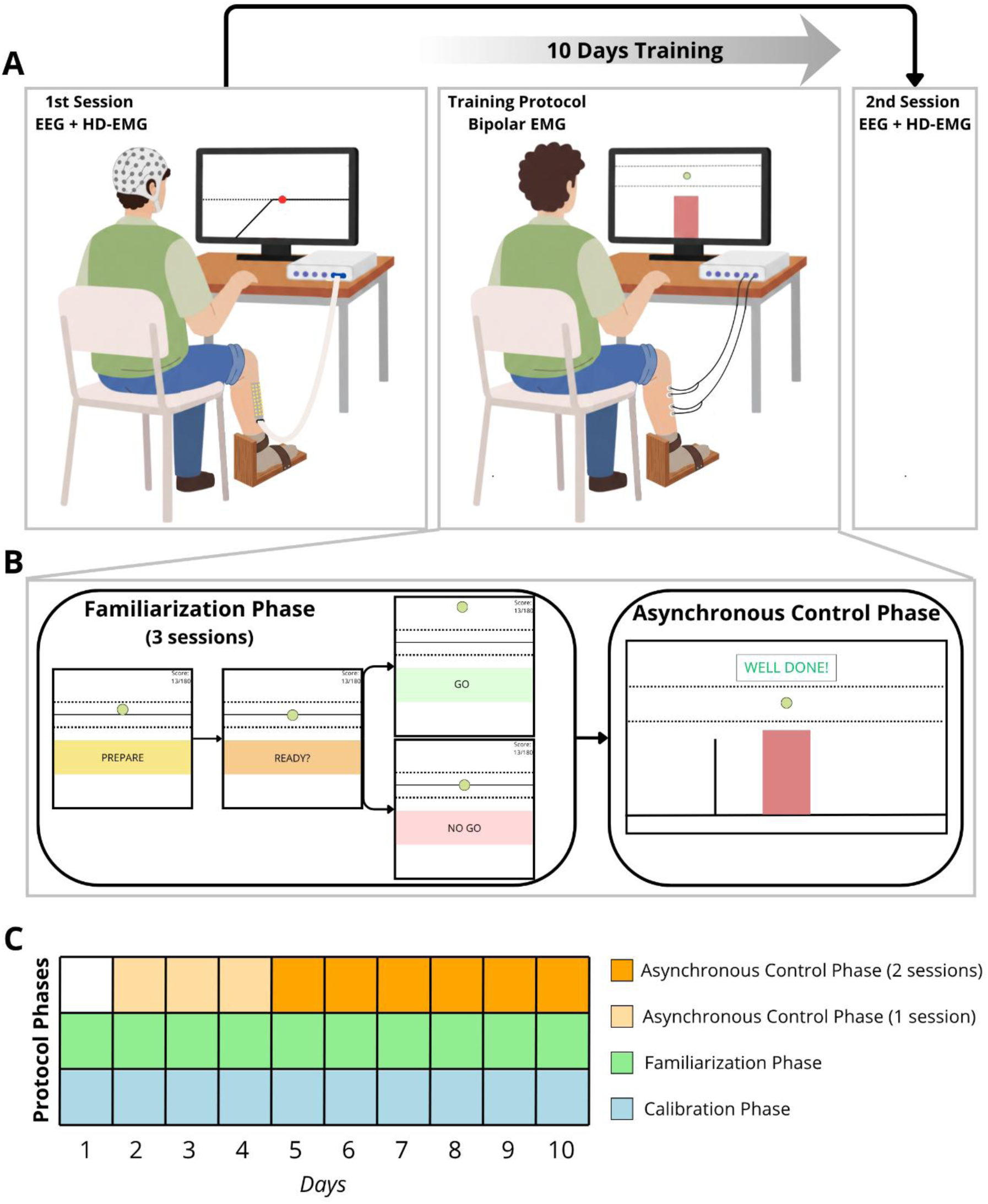
Experimental protocol and neurofeedback-based beta band training paradigm. (A) Overview of the longitudinal experimental design. Participants attended an initial laboratory session in which EEG and high-density EMG (HD-EMG) were recorded while performing a 10% maximum voluntary isometric contraction. This was followed by a 10-day training protocol consisting of repeated neurofeedback sessions, after which participants returned for a final laboratory session with concurrent EEG and HD-EMG recordings. During the 10 days training protocol, participants performed ankle dorsiflexion tasks while receiving real-time visual feedback related to peripheral beta band activity extracted from EMG signals. (B) Structure of the neurofeedback training paradigm. During the familiarization phase (three sessions), participants learned the movement-cancellation strategy used to increase peripheral beta band activity through a GO/NO-GO paradigm. During the subsequent asynchronous control phase, participants practiced voluntarily modulating peripheral beta band activity using a gamified neurofeedback interface. Successful increases in beta band power within the predefined 2-second target window (red shaded area), while the vertical cue remained inside the controlled task region, were rewarded with one point, allowing participants to progressively increase their score. (C) Each training session consisted of a calibration phase to determine the force offset, Maximum Voluntary Contraction, and EMG filter parameters required for neurofeedback generation (see Methods), followed by a familiarization phase. Sessions 1–4 concluded with one asynchronous control block (15 trials) at the end, whereas sessions 5–10 concluded with two asynchronous control blocks (15 trials each).

### Longitudinal Training

Participants were seated comfortably with the knee flexed to ~75° and the leg secured to an ankle dynamometer using straps. The foot rested on a pedal tilted 30° toward plantarflexion (0° representing a neutral ankle angle). Two bipolar EMG electrode pairs were placed over the tibialis anterior as described in the Data Acquisition section. Participants completed neurofeedback-based training sessions using a customized digital interface. Visual information was continuously provided via a monitor placed in front of them. Each participant underwent ten training sessions, scheduled once per day with no more than two days between consecutive sessions. Each session followed the same structured sequence of phases in the order described below. The training protocol is shown in Figure 1C.

#### Calibration phase

This phase opened each session and was used to determine the subject-specific parameters needed for real-time processing. It included force-offset correction, estimation of maximal voluntary contraction (MVC), and a standardized trapezoidal isometric dorsiflexion at 10% MVC. This profile consisted of: a 3-second rest at 0% MVC, a 5-second ramp up to 10% MVC, a 60-second plateau at 10% MVC and a 5-second ramp down to 0% MVC. Intramuscular coherence between the two bipolar EMG channels was computed during the plateau, and the peak beta band coherence frequency was used to define a personalized 5-Hz band-pass filter for real-time beta band feedback as similarly done in Bräcklein et al., (2021). Coherence was estimated using 1-s signal segments and multitaper spectral estimation (three tapers). The upper 95% confidence limit for significance was determined as 1 − 0.05^1/(*L*−1)^, where *L* is the number of segments used in the analysis (Rosenberg et al., 1989).

#### Familiarization phase

This phase introduced the participants to the movement-cancellation mechanism. This mechanism was recently described by Zicher et al., (2024), who showed that cancelling a prepared movement during an isometric contraction elicits a rebound in peripheral beta band power. The rebound was defined as the transient increase in peripheral beta band activity observed following the successful cancellation of a prepared movement. Following the task structure of Zicher et al., (2024), participants performed a GO/NO-GO task consisting of rapid ballistic dorsiflexion on GO cues and inhibition of the prepared movement on NO-GO cues while maintaining a steady 10% MVC baseline (Figure 1B). Each trial consisted of a 5-s preparation period during which participants maintained a 10% MVC dorsiflexion, followed by a ready cue indicating that a response would soon be required. Participants were then presented 1s later with either a GO or NO-GO cue (randomized across trials). During GO trials, participants performed a rapid ballistic dorsiflexion, whereas during NO-GO trials they maintained the 10% MVC contraction. Every trial ended with a 6-s rest period during which force returned to baseline. Each day, this phase consisted of three sessions, each comprising 36 randomized cues (18 GO and 18 NO GO). A 2-minute rest was provided between sessions. The task was gamified using a scoring system to enhance engagement and reinforce correct motor responses. Points could be earned only during GO trials, whereas points could be lost during NO-GO trials if participants produced a movement. Scores were based exclusively on reaction time and force level on each repetition. This scoring system was designed to encourage participants to prepare for movement on every trial, rather than anticipate a NO-GO cue, thereby maximizing engagement of the neural processes associated with the increase in peripheral beta band activity. Reaction time was defined as the interval between the GO cue and the moment force exceeded 14% MVC. Fast and correct responses (<0.55 s) earned +10 points, whereas slower correct responses earned +2 points. NO-GO trials were considered valid only if force remained within 7–13% MVC and was sufficiently stable, with a force SD below 0.5% MVC, as similarly done by Zicher et al. (2024). Any movement during a NO-GO cue resulted in a − 3 point penalty. The maximum score was 180. After each repetition, participants were shown both the score for that trial and their cumulative score on the monitor.

#### Asynchronous control phase

This phase was designed to foster voluntary modulation of beta band activity linked to movement cancellation. Unlike the cue-driven structure of the Familiarization phase, this phase required participants to internally generate and suppress the prepared dorsiflexion movement at precisely timed moments. This phase was introduced progressively from Day 2, starting with one session per day and increasing to two sessions from Day 5 onward to increase the number of trials and overall training load. Each session consisted of 15 trials, during which subjects attempted to voluntarily increase peripheral beta band power.

At the start of this phase, subject-specific parameters from the Calibration and Familiarization phases (MVC, beta band filter width, force offset, EMG data) were loaded to allow real-time force normalization, beta band decoding, and daily beta band threshold computation from valid NO GO trials from the familiarization phase (see Data Analysis). Participants maintained an isometric dorsiflexion at 10% MVC while monitoring force feedback on screen. After a brief stabilization period, 15 events appeared individually at random times on a horizontal timeline. These events were represented as vertical lines moving from right to left toward a fixed 2-s red window (Figure 1B). When an event entered the red window, participants were required to inhibit a prepared movement. Beta band power was computed from a 2-second EMG segment aligned to this inhibition window and compared against the participant-specific daily threshold. Trials in which beta band power exceeded the predefined threshold triggered a positive visual cue and awarded one point, and participants were encouraged to maximize their total score across the 15 trials of each session.

This phase provided participants with direct feedback based on their beta band modulation and marked the transition from externally guided behavioural training to explicit neural control. Trials that failed to meet the predefined force requirements were classified as invalid and no points were awarded.

### Pre- and Post-Training Corticospinal Assessment

These sessions were performed on the first and last day of the protocol to assess whether the 10-day beta-modulation training induced changes in the corticospinal transmission of beta band. During both sessions, we simultaneously recorded cortical activity using EEG, HD-EMG from the tibialis anterior of the dominant leg, and ankle dorsiflexion force.

Similarly to the longitudinal training, participants were seated with the knee flexed at ~75° and the foot secured to an ankle dynamometer positioned at a 30° plantarflexion angle. EEG was acquired using a 31-channel active gel-based cap placed according to the international 10–20 system. A 256-channel grid was placed over the tibialis anterior muscle aligned to the muscle fibers after skin preparation.

Participants then completed four isometric dorsiflexion contractions at different intensities: 5%, 10%, 20%, and 30% of their MVC. Each trial followed a standardized trapezoidal force profile consisting of a ramp-up at 2% MVC per second, a 60-second plateau at the target contraction level, and a ramp-down to rest. A 2-minute break was provided between trials to limit fatigue. The 10% MVC condition matched the force level used during training.

These pre- and post-training sessions provided the data required to evaluate potential training-related changes in beta band corticospinal coupling, neural drive characteristics, and muscle activation patterns.

### Data Analysis

#### Force Analysis

For the familiarization and asynchronous control phases, only NO-GO trials that passed predefined quality criteria were retained for analysis, following an approach similar to Zicher et al., (2024). Force signals were low-pass filtered at 15 Hz with a fourth-order Butterworth filter, baseline-corrected, and normalized to each participant’s MVC. Trials were included only when the force during the NO-GO window remained within 7–13% MVC. Additional rejection criteria were applied to ensure stable isometric contractions: trials were discarded if force deviated by more than ±2% MVC from the average across all accepted trials or if the force standard deviation exceeded 0.5% MVC.

GO trials were considered valid if the force trace exceeded 14% MVC. Reaction time was defined as the interval between the GO cue and the first time point at which the force trace exceeded the 14% MVC threshold.

The coefficient of variation (CoV) of the force was also calculated for the valid windows in which movement was inhibited, as the standard deviation of the force signal divided by its mean value.

#### Beta band features extraction from bipolar EMG

To analyse beta band features, a single bipolar EMG signal obtained from a pair of electrodes positioned over the belly of the tibialis anterior was used for all processing steps. Event markers identified the moment in which participants were required to activate the cancellation mechanism, corresponding to the NO-GO cue during the familiarization phase and to the vertical line entering the inhibition window during the asynchronous control phase. Across days, 56 trials were collected per session in the familiarization phase and between 15 and 30 trials per session in the asynchronous control phase; however, only trials that passed the force-based quality checks (see Force Analysis) were retained. Invalid repetitions, such as performing a GO response during a NO-GO cue, releasing the force prematurely, or failing to maintain a stable 10% MVC contraction, were discarded. This ensured that beta band estimates were derived from consistent and physiologically stable contractions. All retained EMG segments were additionally inspected visually.

EMG was filtered using a subject-specific 5-Hz band-pass fourth-order Butterworth filter centred on the individual peak intramuscular coherence frequency (see Spectral Analysis) identified during the Calibration phase at the beginning of every training day. After filtering, the EMG was squared to obtain its instantaneous power, and the envelope of this activity capturing the time-varying modulation of beta activity was generated. The resulting beta envelope served as the primary signal for all subsequent analyses.

A participant- and day-specific beta threshold was then calculated to determine whether the beta band activity generated during the asynchronous control phase reflected successful activation of the movement-cancellation mechanism. This threshold was derived from the valid NO-GO trials collected in the familiarization phase on that day, under the rationale that these trials require inhibition of a prepared movement and therefore represent the desired neural state. Beta envelope values from all valid NO-GO segments were pooled, and the 98^th^ percentile of this distribution was selected as the daily threshold (Bräcklein et al., 2021). This percentile was chosen to represent a conservative upper limit of naturally occurring beta modulation during successful inhibition while minimising the influence of artifacts. This individualized threshold was then used to classify each trial of the asynchronous control phase as successful or unsuccessful.

In the familiarization phase, temporal and spectral features of beta band activity were extracted from valid NO-GO trials over a 2-s window after the imperative cue. Spectral features included the peak power spectral density (PSD) within the delta (1–4 Hz), theta (4–8 Hz), alpha (8–13 Hz), and beta (13–30 Hz) frequency bands. Temporal features instead characterized the occurrence and evolution of the increase in beta band power in the envelope following movement cancellation, referred to here as the beta band rebound. The beta band rebound was defined as the transient post-cue increase in the peripheral beta band envelope during successful movement cancellation. From this signal, we extracted the beta rebound delay, defined as the latency between the imperative cue and the moment the beta envelope crossed the threshold, as well as the total time for which the envelope remained above threshold.

In the asynchronous control phase, the PSD of the delta, theta, alpha, and beta frequency bands was computed over the 2-s target window for trials in which force remained within the predefined limits (see Force Analysis).

To assess overall muscle activation during movement cancellation, the root mean square (RMS) of the EMG signal was computed for each valid trial over the 2-s window immediately following the NO-GO cue. Only trials satisfying the same force-based inclusion criteria used for the spectral analysis were included. RMS values were pooled across the three sessions of each training day, and the average RMS across valid trials was used as the representative value for each subject and day.

#### Motor Unit decomposition and processing

HD-EMG signals collected before and after the training were decomposed offline to extract individual motor unit discharge patterns using a convolutive blind source separation approach (Negro et al., 2016). As an initial quality check, motor unit spike trains with a silhouette value below 0.9 were automatically excluded (Holobar et al., 2014; Negro et al., 2016). The spike trains that passed this automatic screening were then manually inspected following established procedures (Del Vecchio et al., 2020). During this step, missed firings, typically undetected by peak-detection algorithms (Holobar and Zazula, 2007a, 2007b; Negro et al., 2016), were added, while false detections that produced unrealistic firing rate fluctuations during isometric contractions were removed. This expert-guided correction step increased the accuracy of the motor unit identification used in the subsequent processing. After manual editing, motor unit separation filters were recalculated and re-applied to the raw EMG to refine the final spike train estimates.

Additional quality-control criteria were then applied. Motor units were discarded if their active firing duration covered less than half of the isometric contraction duration, or if their instantaneous firing rate standard deviation exceeded 20 Hz (Del Vecchio et al., 2020). All decomposition steps and manual edits were performed using the MUedit MATLAB interface (Avrillon et al., 2024). Only motor units fulfilling all requirements were kept for further analyses. Each validated motor unit spike train was represented as a binary vector, with ones marking discharge events. Summing these spike trains produced a Cumulative Spike Train (CST), which reflects the common synaptic input received by the motor neuron pool (Farina and Negro, 2015). To ensure CSTs were comparable across conditions, despite differences in the number and firing behaviour of the contributing motor units, we equalized the amplification of the common input across contraction levels. Because the CST’s gain scales with the number of motor unit discharges (determined by MU recruitment and firing rate; (Farina et al., 2014)), we first identified the contraction with the lowest total firing count and then randomly subsampled motor units from the remaining conditions to match this firing count. This process was repeated 10 times for each contraction level, generating 10 CST permutations. All subsequent analyses were performed on these CSTs, and final values for each participant and condition were taken as the average across all permutations. This approach has previously been used in similar analyses (Abbagnano et al., 2025).

Subject 6 was excluded from all analyses involving decomposed motor units due to technical issues that prevented reliable HD-EMG recordings for that participant.

#### EEG processing

The signals were first re-referenced offline using the average of the earlobe electrodes collected during the recording. A fourth-order Butterworth band-pass filter (0.5–45 Hz) was then applied. Artifact components associated with blinks, eye movements and muscular activity were identified and removed using Independent Component Analysis. To further improve spatial specificity, a surface Laplacian transform was computed with the CSD Toolbox (Kayser and Tenke, 2006), reducing shared activity across nearby electrodes. The EEG preprocessing pipeline was carried out in FieldTrip (Oostenveld et al., 2011).

The ‘Cz’ electrode was used for the analysis because it typically shows the strongest corticomuscular coherence in the beta band during tibialis anterior contractions (Ibáñez et al., 2021).

#### Spectral Analysis

Spectral analyses were performed to characterise the frequency-domain properties of cortical and peripheral neural activity across the different phases of the protocol. For the familiarization and asynchronous-control phases, the PSD of the bipolar EMG was computed during valid NO-GO windows to quantify power distribution across frequencies in different bands. PSD estimates were obtained using Welch’s method in MATLAB (pwelch, 1-second windows, 50% overlap). The peak PSD within each frequency band and its corresponding peak frequency were then extracted. The same procedure was applied to the EEG and HD-sEMG recordings collected before and after the training to evaluate beta band activity at both cortical and peripheral levels. In these sessions, PSD was computed using Welch’s method with 1-second windows and 50% overlap.

Coherence analyses were carried out with the Neurospec 2.11 toolbox (Halliday, 2015). Corticomuscular coherence between the decomposed cumulative spike train and the Cz channel was used to assess differences in the transmission of cortical beta oscillations to the tibialis anterior before and after training. Intramuscular coherence was also computed daily between the two bipolar EMG channels during the calibration phase to quantify the common synaptic input to the motor neuron pool. Intramuscular coherence was calculated over a stable 50-second segment of isometric contraction at 10% MVC, starting two seconds after reaching the target force to avoid transient effects. Before estimating coherence, both EMG signals were rectified and detrended. For each participant, training session, and coherence measure, the maximum coherence value within the beta band (13-30 Hz) was selected for analysis.

#### NASA-TLX Analysis

Subjective workload was evaluated starting from the second day of the training at the end of each session using the NASA-TLX questionnaire (Hertzum, 2021). This measure was used to track whether participants perceived the task as becoming less demanding over time, reflecting growing familiarity and efficiency. We focused in particular on changes in the Effort, Physical Effort, and Mental Effort subscales, as these dimensions are most relevant to sustained isometric control and beta band modulation.

#### Statistical Analysis

All statistical analyses were performed using Jamovi (www.jamovi.org) and custom MATLAB scripts. Data are reported as mean ± standard deviation (SD), and statistical significance was set at p < 0.05. The Shapiro–Wilk test was used to evaluate whether each variable followed a normal distribution.

Longitudinal changes across the 10-day training protocol were assessed using Linear Mixed Models (LMMs). For each dependent variable, an LMM was constructed with *day* entered as a fixed effect and *participant* included as a random intercept to account for repeated-measures within subjects. The Fixed Effects Omnibus Test was used to determine whether *day* had a significant overall influence on the variable of interest. For all models showing a significant effect (p < 0.05), the corresponding F-values are reported.

To compare spectral measures, including PSD peaks from EEG and CST signals as well as corticomuscular coherence, before and after training, we used a non-parametric repeated-measures test (Friedman test), as these variables did not satisfy normality assumptions.

## Results

We tested whether beta oscillations can be voluntarily controlled through training by exploiting the characteristic increase in beta power that occurs after movement cancellation, illustrated by the representative beta band envelopes shown in Figure 2. We also tested whether these behavioural changes were accompanied by adaptations in both peripheral and cortical beta band activity. Participants completed a 10-day neurofeedback protocol comprising a guided familiarization phase and an asynchronous control phase, both aimed at learning and voluntarily engaging the movement-cancellation mechanism to increase peripheral beta band power. We then assessed behavioural performance, peripheral beta band activity, and corticospinal transmission using concurrent EEG and HD-EMG recordings collected before and after training.

**Figure 2.**
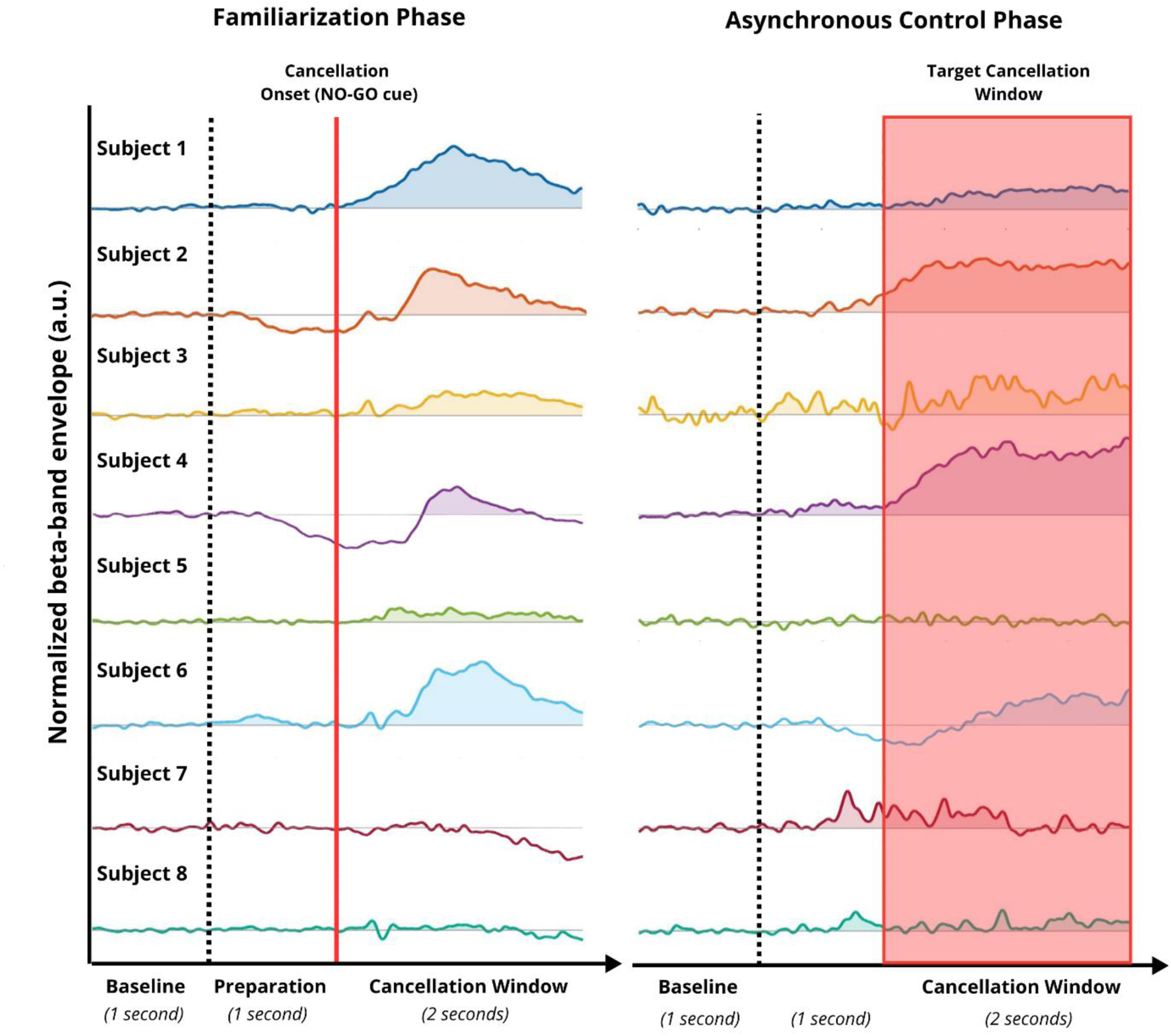
Representative beta band EMG envelopes during movement cancellation. Traces represent participant-specific beta band EMG envelopes averaged across repetitions and days in the familiarization (left) and asynchronous control (right) phases. EMG signals were band-pass filtered between 13 and 30 Hz, converted to amplitude envelopes using the Hilbert transform, and baseline-corrected. Colored traces identify individual participants, and shaded regions indicate positive deviations from baseline. The black dashed line denotes movement-preparation onset, whereas the red line indicates cancellation onset. In the familiarization phase, movement cancellation was externally cued; in the asynchronous control phase, participants voluntarily suppressed a prepared movement when the feedback-controlled cursor entered the 2-s target window (see Methods).

### Enhanced beta band during the cancellation period

To verify that participants performed the task correctly, we quantified the percentage of valid trials based on the criteria described in the *Methods* section. The percentage of valid NO-GO trials across the 10 training days is reported in Table 1, together with the delay between the GO cue and the time at which force exceeded 14% MVC, the threshold used to define a valid GO trial. The force traces for the valid GO and NO-GO trials are reported in Figure 3A.

**Figure 3.**
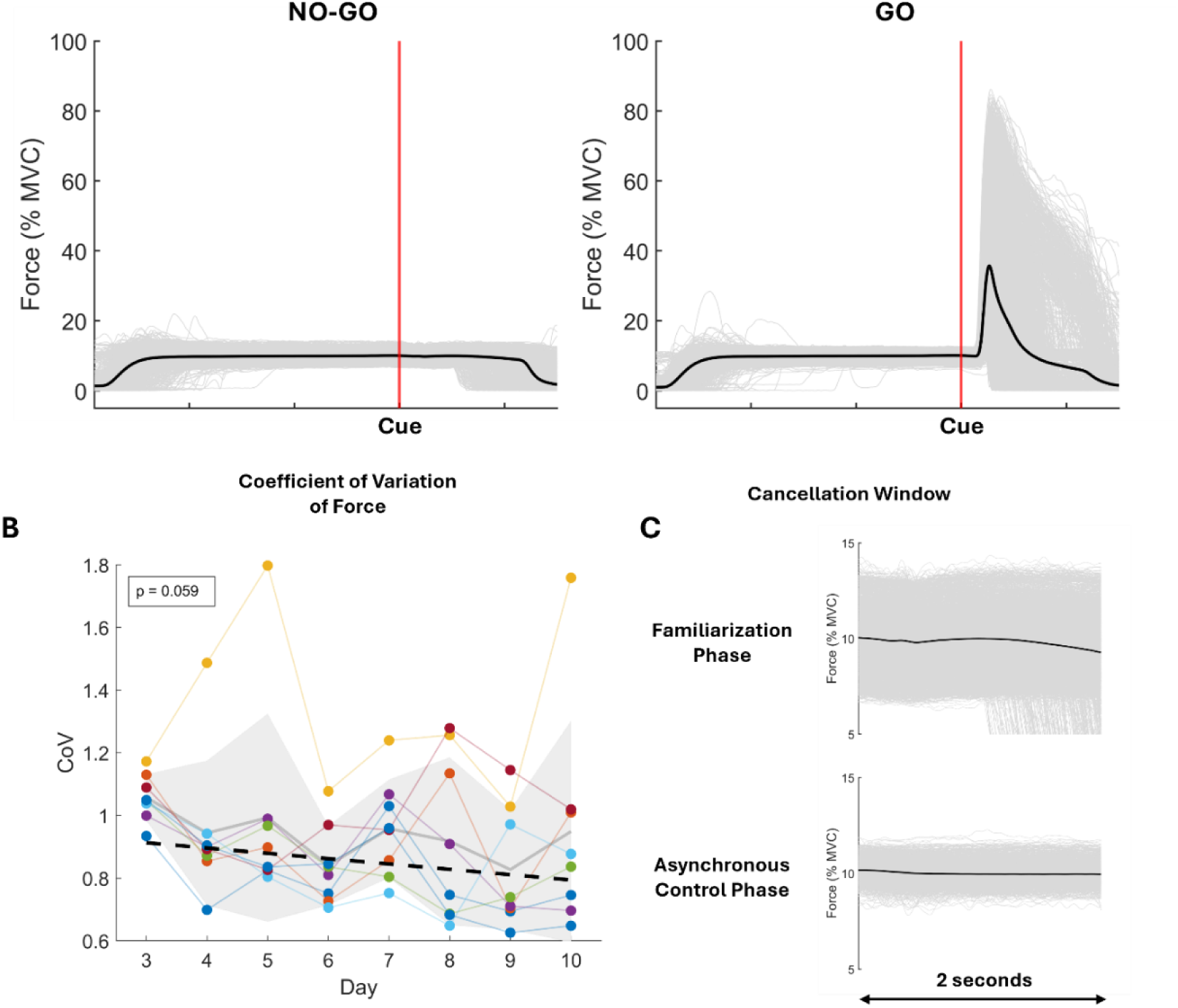
Force analysis during the training protocol. Representative force traces from correctly performed GO and NO-GO trials during the Familiarization phase are shown in A. Individual trials are displayed in light gray, with the average across all trials shown in bold black. The red vertical line indicates the onset of the GO or NO-GO cue following the preparation period. The coefficient of variation (CoV) of force during the cancellation window across training days is shown in B. Each point represents the average measurement for a given day. Daily values are normalised to the mean of the first two training days, which are therefore not displayed. The black dashed line indicates the fixed-effect linear trend estimated by the Linear Mixed Model. Individual participants are shown in distinct colours. The light gray line represents the group mean, and the shaded gray region denotes ±1 standard deviation across participants. In C, individual force traces (light gray) and the average trace (black) are shown during the 2-s cancellation window following the NO-GO cue in the Familiarization phase, and during the corresponding period in the Asynchronous Control phase, when participants were instructed to voluntarily increase beta band power by engaging the learned cancellation mechanism (see Methods).

**Table 1.**
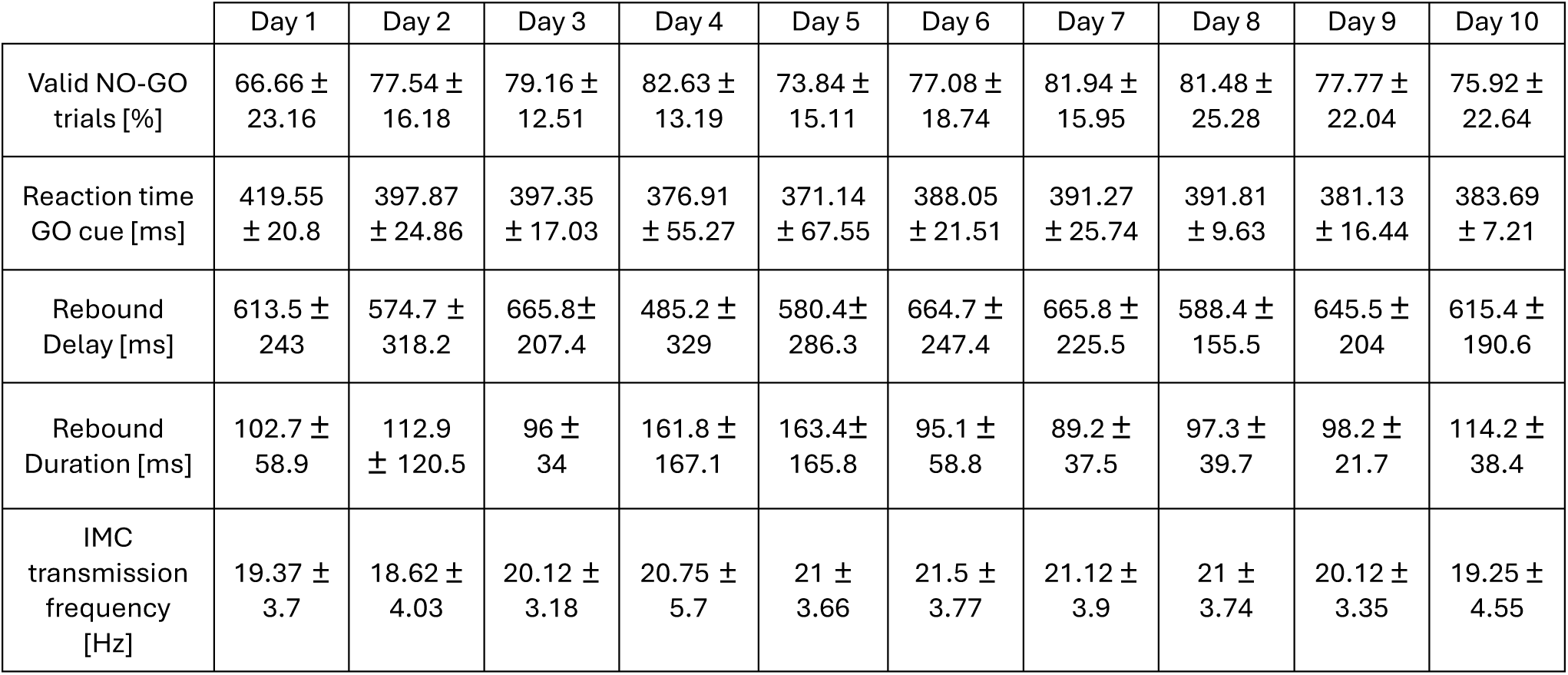
Summary of experimental results. Mean ± standard deviation of experimental measures across the 10 days of the training protocol. Valid NO-GO trials refer to trials performed during the familiarization phase. Rebound refers to the increase in beta band power observed during movement cancellation in the NO-GO condition of the familiarization phase. Rebound delay and duration are expressed in milliseconds and were quantified relative to a participant-specific threshold defined as the 98th percentile of the beta band envelope. Delay corresponds to the time elapsed before the envelope exceeded the threshold after the NO-GO cue, whereas duration corresponds to the time the envelope remained above it. IMC transmission frequency refers to the frequency (Hz) at which the intramuscular coherence (IMC) spectrum, estimated from bipolar EMG signals recorded during the calibration phase at the beginning of each training session, exhibited its peak value.

|  | Day 1 | Day 2 | Day 3 | Day 4 | Day 5 | Day 6 | Day 7 | Day 8 | Day 9 | Day 10 |
| --- | --- | --- | --- | --- | --- | --- | --- | --- | --- | --- |
| Valid NO-GO trials [%] | 66.66 ± 23.16 | 77.54 ± 16.18 | 79.16 ± 12.51 | 82.63 ± 13.19 | 73.84 ± 15.11 | 77.08 ± 18.74 | 81.94 ± 15.95 | 81.48 ± 25.28 | 77.77 ± 22.04 | 75.92 ± 22.64 |
| Reaction time GO cue [ms] | 419.55 ± 20.8 | 397.87 ± 24.86 | 397.35 ± 17.03 | 376.91 ± 55.27 | 371.14 ± 67.55 | 388.05 ± 21.51 | 391.27 ± 25.74 | 391.81 ± 9.63 | 381.13 ± 16.44 | 383.69 ± 7.21 |
| Rebound Delay [ms] | 613.5 ± 243 | 574.7 ± 318.2 | 665.8 ± 207.4 | 485.2 ± 329 | 580.4 ± 286.3 | 664.7 ± 247.4 | 665.8 ± 225.5 | 588.4 ± 155.5 | 645.5 ± 204 | 615.4 ± 190.6 |
| Rebound Duration [ms] | 102.7 ± 58.9 | 112.9 ± 120.5 | 96 ± 34 | 161.8 ± 167.1 | 163.4 ± 165.8 | 95.1 ± 58.8 | 89.2 ± 37.5 | 97.3 ± 39.7 | 98.2 ± 21.7 | 114.2 ± 38.4 |
| IMC transmission frequency [Hz] | 19.37 ± 3.7 | 18.62 ± 4.03 | 20.12 ± 3.18 | 20.75 ± 5.7 | 21 ± 3.66 | 21.5 ± 3.77 | 21.12 ± 3.9 | 21 ± 3.74 | 20.12 ± 3.35 | 19.25 ± 4.55 |

Repeating the familiarization phase across days was intended to strengthen subjects’ understanding of movement cancellation, a process known to coincide with an increase in beta band power. Beyond behavioural practice, repeated exposure could induce physiological adaptations that may influence how easily this signal can be modulated later.

To test for such changes, we analysed the PSD in the beta band during valid NO-GO trials from day 3 to day 10, normalized to the mean power of the first two days that were thus used as baseline. We observed a significant increase in beta band power across days (Figure 4D; F(1,55) = 4.64; p = 0.036). This increase varied across participants, with some showing a clearer trend than others. Notably, this increase was specific to the beta band, as no significant changes were observed in the delta (Figure 4A; F(1,55) = 1.26; p = 0.267), theta (Figure 4B; F(1,55) = 1.77; p = 0.188), or alpha bands (Figure 4C; F(1,55) = 3.54; p = 0.065). No significant change in the EMG RMS was observed over the same analysis window (F(1,55) = 2.54, p = 0.117). Overall, these results indicate that the familiarization phase was associated with stronger peripheral beta band power after movement cancellation onset, possibly reflecting greater synchronization of motor unit activity specifically in the beta band.

**Figure 4.**
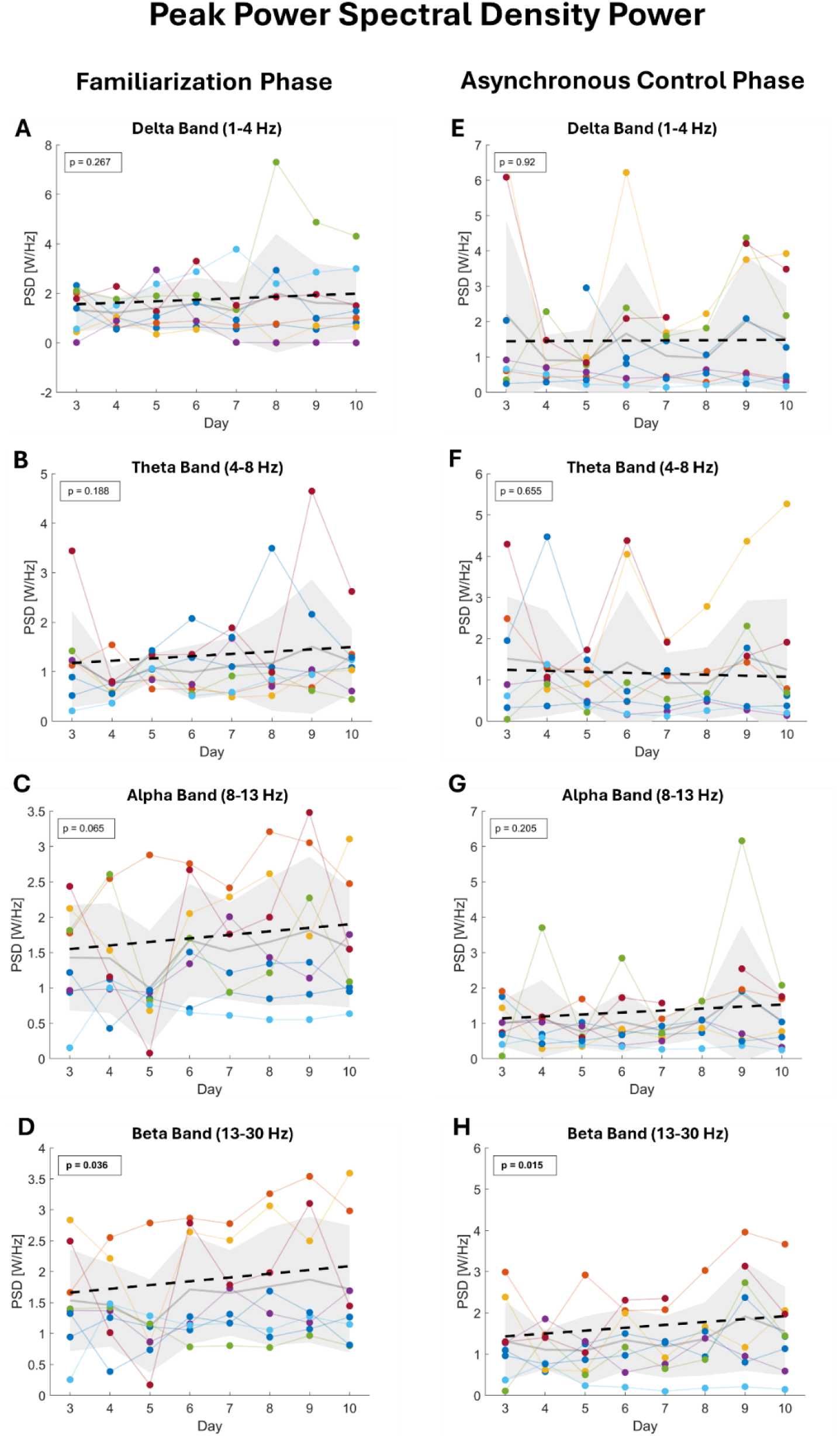
Evolution of power spectral density (PSD) peak values across the training protocol days. Average peak power spectral density (PSD) across training days for the Familiarization phase (left; A–D) and Asynchronous control phase (right; E–H) in the delta (1–4 Hz; A,E), theta (4–8 Hz; B,F), alpha (8–13 Hz; C,G), and beta (13–30 Hz; D,H) frequency bands. Individual participants are shown in distinct colours. Daily peak PSD values were normalized to the mean of the first two training days, which are therefore not displayed. Each point represents the average measurement for a given day. The black dashed line indicates the fixed-effect linear trend estimated by the Linear Mixed Model. The light gray line represents the group mean, and the shaded gray region denotes ±1 standard deviation across participants.

We also examined whether the temporal characteristics of this beta band rebound, defined here as the post-cue increase in the peripheral beta band envelope during successful movement cancellation, changed with training. Specifically, we analysed the delay between the imperative NO-GO cue and the moment when the beta envelope exceeded the daily threshold, as well as the duration of the rebound above threshold. Neither the delay (F(1,64.2) = 1.126; p = 0.723) nor the rebound duration (F(1,65.9) = 0.175; p = 0.677) changed significantly across training days (see Table 1). The threshold was defined as the 98th percentile of the beta band envelope values computed from the valid NO-GO trials (see Methods).

Taken together, these findings show that the only systematic training-related change during the familiarization phase was an increase in beta band power during movement cancellation, whereas the timing and duration of the beta rebound remained stable. This result is consistent with previous evidence suggesting that beta band oscillations are transmitted through the fastest corticospinal pathways, potentially limiting the extent to which their temporal characteristics can be modified (Ibáñez et al., 2021).

Next, we examined whether the increase in beta band power observed during the cancellation window in the familiarization phase was also present during the asynchronous control phase, given that both phases engage the same underlying neural mechanism. Although beta band power was computed in the same way in the two phases, the tasks differed markedly: the familiarization phase followed a guided GO/NO-GO structure with cues, whereas the asynchronous control phase required self-initiated, unguided modulation of beta activity in a specific target window. These differences in cognitive and motor demands could influence how strongly beta band modulation emerges during training. Despite these task-level distinctions, we observed a comparable increase in beta band power across days in the asynchronous control phase (Figure 4H; F(1,54) = 6.26; p = 0.015). Although the magnitude of this increase varied across participants, the effect was specific to the beta band. No significant training-related changes were detected in the delta (Figure 4E; F(1,53.1) = 0.01; p = 0.92), theta (Figure 4F; F(1,53) = 0.2; p = 0.655), or alpha (Figure 4G; F(1,54.1) = 1.65; p = 0.205) bands. Similarly to the familiarization phase, no significant changes in EMG RMS were observed across training days (F(1,54) = 2.17; p = 0.146).

Given the observed increase in beta band power, we examined whether this was accompanied by a corresponding rise in the relative strength of common synaptic input to the motor neuron pool in the beta range. Intramuscular coherence was computed daily using the two bipolar EMG signals acquired during the calibration phase while participants performed a 10% MVC isometric contraction (see Methods). There was a progressive increase in the intramuscular coherence spectrum across the training days relative to baseline, which was defined as the average intramuscular coherence measured during the first two training days to reduce variability (Figure 5A). Across the 10 days, intramuscular coherence in the beta band increased significantly (Figure 5B; F(1,55) = 6.08; p = 0.017) indicating that the increase in beta band power was accompanied by a strengthening of the common input to the motor neuron pool in this band. We also tested whether the average transmission frequency of this input in the beta band changed across training days (see Table 1). No significant effect was found (F(1,71) = 0.59; p = 0.443), suggesting that while the magnitude of the beta band common input increased, its central frequency remained stable.

**Figure 5.**
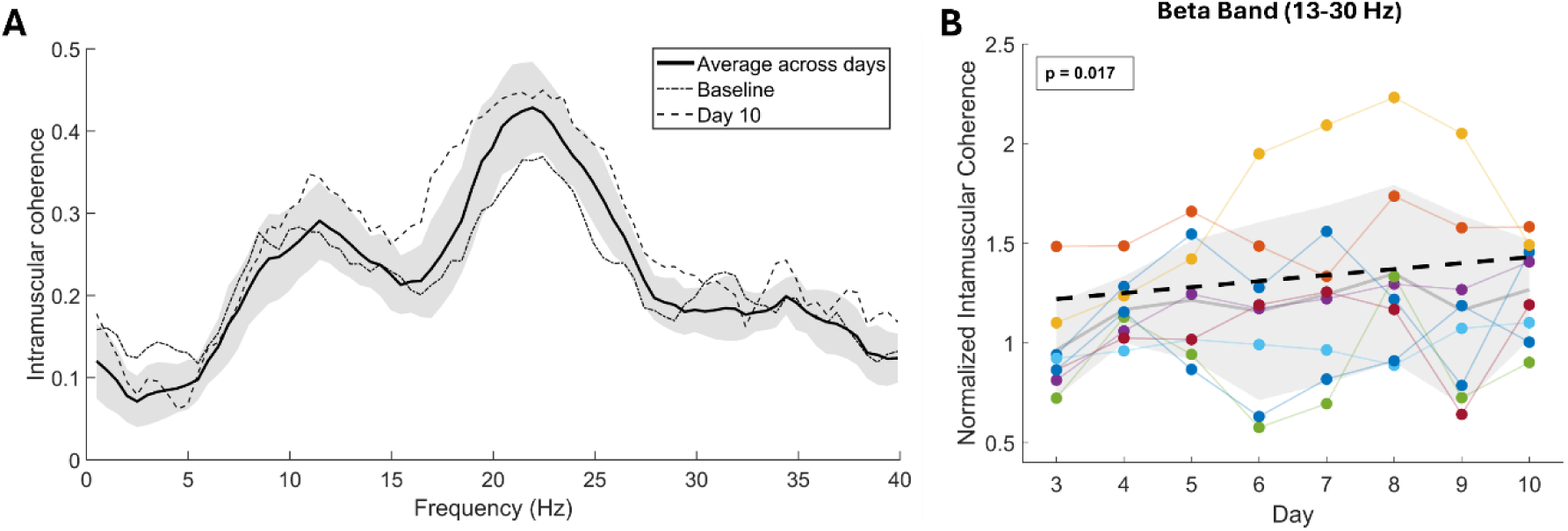
Intramuscular coherence and beta band peak coherence across the training protocol. Intramuscular coherence spectra between 0 and 40 Hz are shown in A. The baseline spectrum, computed as the average of the first two training days, is displayed as a dash-dotted line, while the spectrum from the final training day (day 10) is shown for comparison as a dashed line. The solid black line represents the average coherence spectrum across all training days, and the light gray shaded region denotes ±1 standard deviation. In B, the average peak intramuscular coherence within the beta band is shown across training days. Daily values were normalized to the mean of the first two training days, used as baseline, which are therefore not displayed. Each point represents the average measurement for a given day. The black dashed line indicates the fixed-effect linear trend estimated by the Linear Mixed Model. The light gray line represents the group mean, and the shaded gray region denotes ±1 standard deviation across participants.

Participants showed a trend toward improved force control across the training protocol. Specifically, we observed a trend toward a reduction in the coefficient of variation of force (Figure 3B; F(1,55) = 3.73; p = 0.059), suggesting that participants became progressively more stable during movement cancellation over time. Force traces recorded during movement inhibition are shown in Figure 3C. These correspond either to the period immediately following the NO-GO cue during the familiarization phase or to successful voluntary engagement of the movement-cancellation mechanism within the target window during the asynchronous control phase.

### Training enhances voluntary modulation of the beta band signal

While the longitudinal training produced clear changes in beta band activity, both in terms of power and common input, the key question is whether activating this mechanism actually became easier for participants. Because beta band activity was decoded peripherally and no direct functional role has been identified, intentionally modulating it is challenging. The observed increase in beta power associated with a more constrained mental strategy could facilitate its voluntary control, potentially enabling higher scores during the asynchronous control phase. However, because this phase was entirely unguided, improvements depended on each participant’s ability to develop an effective internal strategy that leveraged the movement-cancellation mechanism acquired during the familiarization phase.

Across days, we observed a significant increase in the overall daily score, indicating that participants became increasingly more successful at generating elevated beta band power within the required target window during the asynchronous control phase (Figure 6; F(1,62.1) = 7.3; *p* = 0.009). Specifically, the score reflected the number of successful trials, out of 15 per session, in which the beta rebound exceeded the daily participant-specific threshold within the required target window. Although this trend is evident at the group level, the magnitude of improvement varied substantially across individuals, indicating heterogeneous learning trajectories.

**Figure 6.**
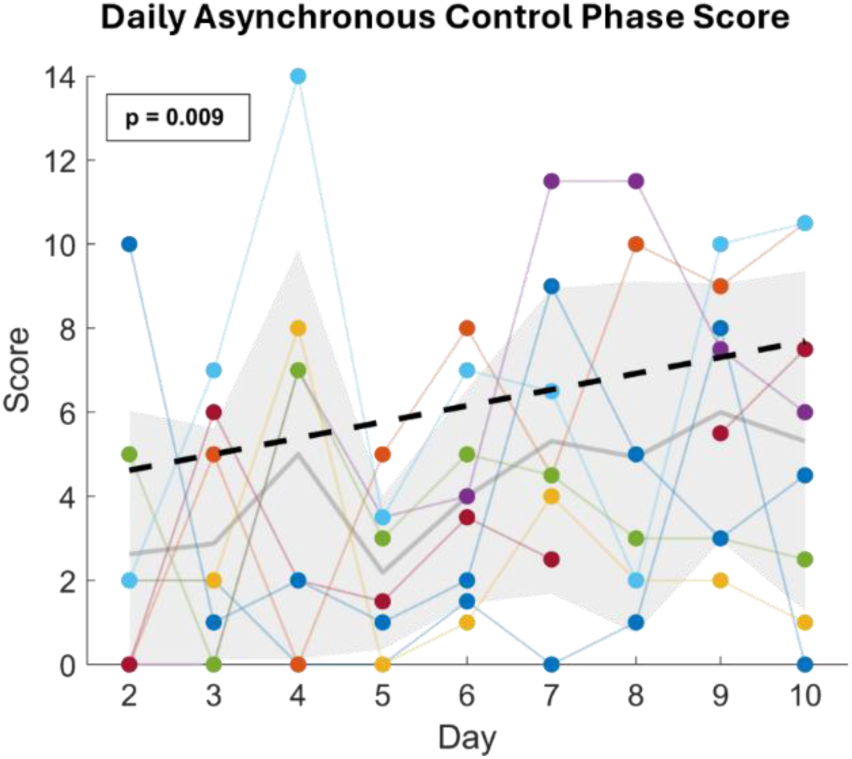
Score achieved during the asynchronous control phase across training days. The score represents the number of successful trials (maximum = 15 per session), with one point awarded each time beta band power exceeded the predefined threshold (see Methods). Because the score was not normalized to the first training day, values are presented starting from Day 2 of the training protocol, which corresponds to the first day of the asynchronous control phase. For each plot, individual subjects are shown in distinct colours. The black dashed line shows the fixed-effect linear trend estimated by the Linear Mixed Model. The light gray line indicates the mean across participants, and the shaded gray region represents the standard deviation.

To evaluate whether this improvement reflected a subjective reduction in task difficulty, we examined workload reports collected through the NASA-TLX questionnaire. Participants reported a significant decrease in the perceived overall effort required to control the interface (Figure 7A; F(1,63) = 6.57; *p* = 0.013). However, neither physical effort (Figure 7B; F(1,63) = 1.87; *p* = 0.177) nor mental effort (Figure 7C; F(1,63) = 1.17; *p* = 0.283) showed significant changes across days. This suggests that while participants became more effective at generating successful trials, the task itself did not feel easier from a physical or cognitive standpoint.

**Figure 7.**
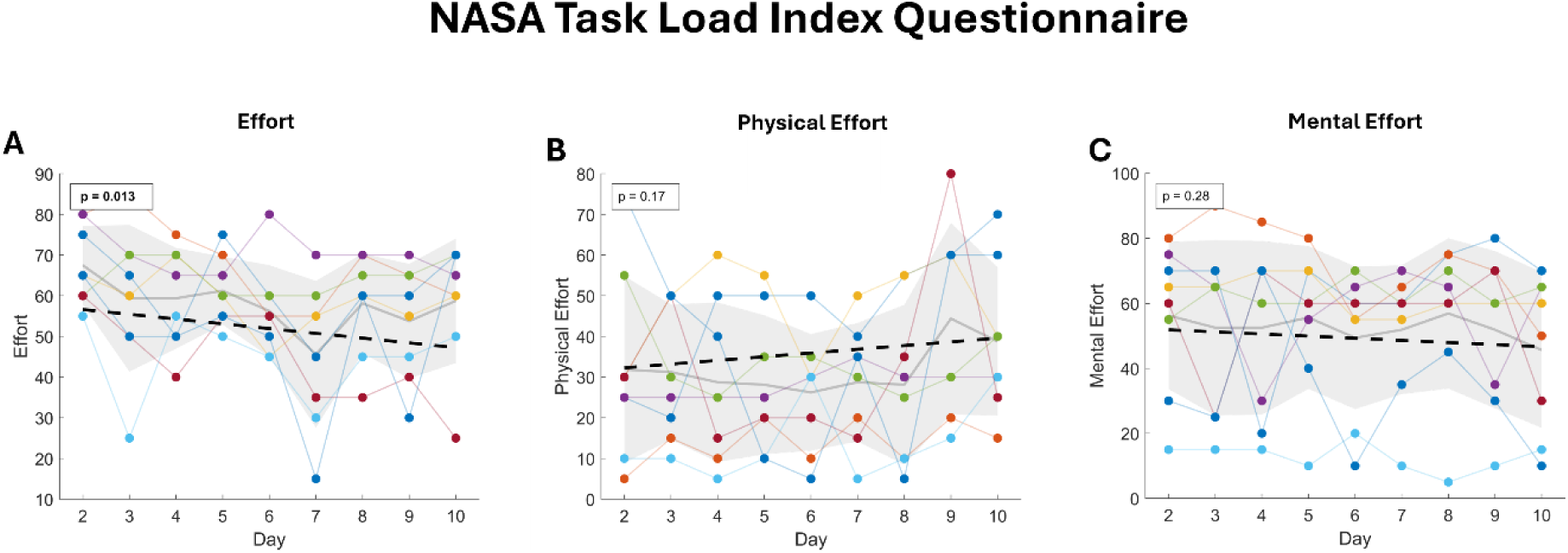
NASA Task Load Index reported values. For each plot, individual subjects are shown in distinct colours. The black dashed line shows the fixed-effect linear trend estimated by the Linear Mixed Model. The light gray line indicates the mean across participants, and the shaded gray region represents the standard deviation.

### No Training-Related changes in the corticospinal transmission

Participants completed dedicated sessions of concurrent EEG and HD-EMG recordings to examine whether corticospinal transmission of beta band activity was altered by the 10-day protocol. These assessments were performed during steady isometric dorsiflexion contractions at 5%, 10%, 20%, and 30% MVC. To obtain a more precise estimate of the common input to motor neurons in the beta band (Day and Hulliger, 2001; Keenan et al., 2006), HD-EMG signals were decomposed into individual motor unit spike trains. The number of valid identified motor units (see Methods) remained comparable before and after training across all force levels: 9.71 ± 5.93 vs.10.85 ± 4.74 at 5% MVC (χ^2^(1) = 0.2; p = 0.655), 16 ± 8.3 vs. 17.14 ± 9.06 at 10% MVC (χ^2^(1) = 0.667; p = 0.414), 13.71 ± 8.3 vs. 14 ± 7.3 at 20% MVC (χ^2^(1) = 0.143; p = 0.705), and 7.42 ± 4.42 vs. 9.14 ± 4.87 at 30% MVC (χ^2^(1) = 1.29; p = 0.257).

We assessed corticomuscular coupling using coherence analysis and found no significant differences before and after training at 10% MVC (χ^2^(1) = 0.143; p = 0.705), 20% MVC (χ^2^(1) = 1.29; p = 0.257), or 30% MVC (χ^2^(1) = 1.29; p = 0.257), with the exception of 5% MVC, where a significant effect was observed (χ^2^(1) = 7; p = 0.008) (Figure 8C). Similarly, cortical beta band power showed no change at any contraction level (5% MVC: χ^2^(1) = 0.143, p = 0.705; 10% MVC: χ^2^(1) = 0.143, p = 0.705; 20% MVC: χ^2^(1) = 0.143, p = 0.705; 30% MVC: χ^2^(1) = 1.29; p = 0.257) (Figure 8B).

**Figure 8.**
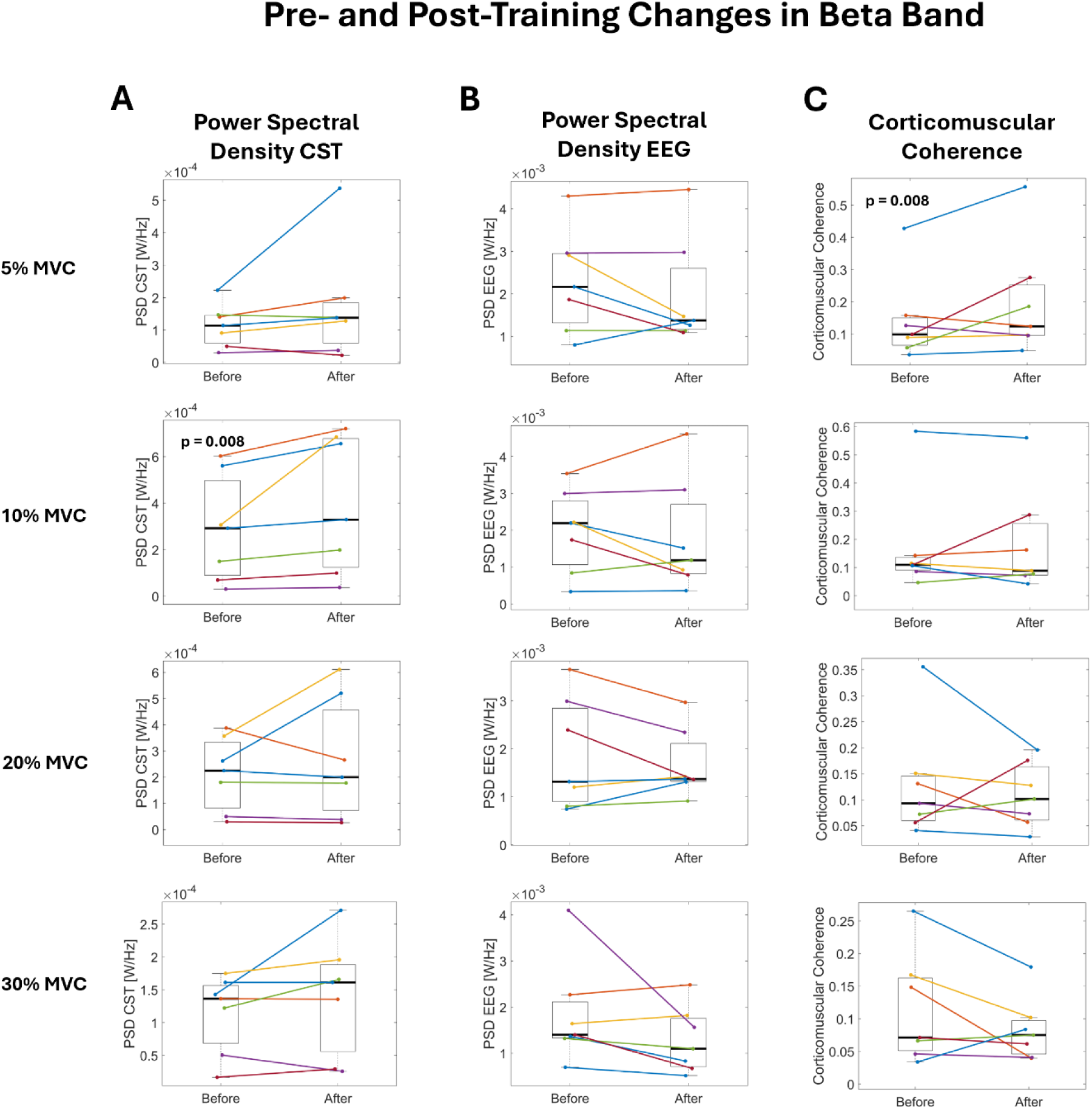
Results from the concurrent EEG and HD-EMG isometric contraction sessions recorded before and after the 10-day training protocol. For each plot, individual subjects are shown in distinct colours. (A) Peak power spectral density (PSD) in the beta band for the Cumulative Spike Train (CST). (B) Peak PSD in the beta band for the ‘Cz’ EEG channel. (C) Corticomuscular coherence between the ‘Cz’ EEG channel and the CST.

In contrast, peripheral beta activity derived from the Cumulative Spike Train increased significantly at 10% MVC (χ^2^(1) = 7; p = 0.008), whereas no changes were observed at 5% (χ^2^(1) = 1.29; p = 0.257), 20% (χ^2^(1) = 1.29; p = 0.257), or 30% MVC (χ^2^(1) = 1.29; p = 0.257) (Figure 8A). This selective enhancement at the training intensity aligns with the increases observed during the cancellation windows and indicates that training-related adaptations occurred predominantly at the spinal or motor unit level, rather than at cortical level.

## Discussion

This study shows that peripheral beta band activity can be strengthened through targeted training. Across a 10-day protocol, participants not only learned to engage the movement-cancellation mechanism more successfully but also demonstrated progressive improvements in their ability to voluntarily increase beta band power during the asynchronous control phase. These behavioural gains were accompanied by physiological adaptations at the peripheral level: beta band power increased during cancellation, and the common synaptic input to the motor neuron pool in this band strengthened across days. These changes occurred without measurable alterations in corticospinal transmission, indicating that the adaptations arose primarily subcortically.

### Better behavioural control emerges alongside strengthened peripheral beta power

We designed a longitudinal training paradigm that leverages the physiological increase in peripheral beta band power that emerges when a prepared movement is intentionally inhibited (Zicher et al., 2024). Overall, participants improved their performance across days, with a significant increase in the group-level score, defined as the number of trials in which participants increased peripheral beta band power above a daily subject-specific threshold. Notably, these improvements were achieved using only bipolar EMG, a signal affected by amplitude cancellation and therefore providing reduced access to detailed neural information compared with motor unit decomposition using HD-EMG (Farina et al., 2008; Keenan et al., 2006).

As participants improved their ability to engage this two-state control signal (i.e., the increase in beta band power associated with cancelling a prepared movement), we also observed clear peripheral adaptations that were largely confined to the beta band and not to other bands. Nonetheless, the changes in alpha band had a clear tendency, although non-significant, to change, which is consistent with previous studies reporting increased alpha band activity during movement preparation and response inhibition in GO/NO-GO paradigms (Pani et al., 2014; Picazio et al., 2014; Zicher et al., 2024). Beta band power increased during movement cancellation not only in the familiarization phase but also in the asynchronous control phase, despite the substantial differences between the two tasks.

This enhancement in peripheral beta activity was not restricted to the brief cancellation window. Participants also performed a sustained isometric contraction at 10% MVC during the daily calibration phase and we examined whether training altered beta band features during this sustained muscle activation. We found a significant increase in the common synaptic input to the motor neuron pool, as indicated by intramuscular coherence. Furthermore, motor unit decomposition showed that beta band power estimated from the cumulative spike train during 10% MVC contractions increased after training compared with before training. This effect was specific to the contraction level used during the training sessions and was not observed at other intensities.

In contrast, no training-related changes emerged at the cortical level: neither EEG beta power nor corticomuscular coherence showed significant modulation (except for a small effect at 5% MVC for the corticomuscular coherence). This selective increase in peripheral beta band power raises questions regarding the factors of influence of peripheral beta. In a previous study (Abbagnano et al., 2025), we reported that the power of beta band oscillations at both the cortical and peripheral levels remained comparable across isometric contractions at increasing force levels. Therefore, if the increase in peripheral beta band activity originated primarily from cortical adaptations a comparable increase would be expected across all contraction intensities rather than being limited to the 10% MVC contraction.

These findings indicate that the physiological adaptations associated with this protocol were mainly peripheral, suggesting that the training primarily influenced spinal level mechanisms rather than corticospinal transmission.

This pattern is consistent with the broader view proposed by Wolpaw & Kamesar (2022), whereby the acquisition of a new control strategy is accompanied by distributed adaptations that extend beyond the specific task variable used for feedback. In our case, the behavioural improvement was not associated solely with greater beta band power, which was the signal directly rewarded during training, but also with changes in more general measures of neural output within the muscle, such as the intramuscular coherence. This suggests that training did not simply enhance performance on the task itself, but may have induced a broader adaptation of the motor system, expressed at least at the level of spinal circuitry and motor neuron output. Although we did not detect cortical changes with EEG, this does not exclude cortical contributions, as subtle training related adaptations may have remained below the resolution or signal-to-noise limits of the technique.

### Peripheral mechanisms underlying improved beta control

Peripheral beta band power increased during movement cancellation, and participants also became more stable across days, as indicated by the progressive reduction in the coefficient of variation for force. The association between greater force stability and stronger beta band activity is consistent with the status-quo hypothesis, which proposes that beta activity supports maintenance of the current motor state while opposing transitions to new movements (Engel and Fries, 2010; Gilbertson et al., 2005; Siegel et al., 2009). Within this framework, training may have progressively enhanced participants’ ability to stabilize the ongoing motor state during movement inhibition. While this hypothesis provides a functional interpretation of our findings, the neural mechanisms underlying the increase in peripheral beta band activity remain unclear. One potential spinal mechanism is a training-induced adaptation of inhibitory circuitry, including recurrent inhibition. Recurrent inhibition can shape the frequency content of motor neuron output and interact with common synaptic input in the beta band, thereby influencing the expression of synchronized activity at the motor unit level (Dernoncourt et al., 2025; Williams and Baker, 2009b). Repeated movement cancellation could therefore have modified this inhibitory balance, facilitating the emergence of peripheral beta band activity. However, because recurrent inhibition was not measured directly in the present study, this mechanistic interpretation remains speculative.

### Translational potential of peripheral beta band modulation

Neural oscillations in the beta band are widespread across the motor system and are closely linked to motor control (Baker et al., 1999; Kilavik et al., 2013), although their precise functional role remains debated (Engel and Fries, 2010). Beta band activity also varies substantially across individuals and is altered in several neurological conditions, including Parkinson’s disease, stroke, and spinal cord injury (Gourab and Schmit, 2010; Matsuya et al., 2017; Ushiyama et al., 2011; Von Carlowitz-Ghori et al., 2014; Zokaei et al., 2021). These observations raise the question of whether peripheral beta band activity can be modified through training. While motor learning has been shown to alter cortical beta band activity (Moisello et al., 2015), this had not previously been demonstrated at the peripheral level. Our findings show that peripheral beta band activity is not fixed, but can be shaped through targeted behavioural training. This suggests that beta band plasticity may be relevant not only for understanding motor control, but also for future interventions in neurological populations with altered oscillatory activity.

Additionally, a biological signal that can be both decoded and voluntarily shaped through targeted training represents a valuable candidate control signal for motor augmentation. An effective control signal for augmentation should be independent of natural movements, reliably generated, consistently measurable, and at least partly under voluntary control (Eden et al., 2022). Peripheral beta band activity appears to meet several of these requirements. In addition, our findings show that it is possible to design a mental strategy for its control and that performance can improve with longitudinal training.

However, although participants became more effective at generating successful trials, the task did not necessarily feel easier from a physical or cognitive perspective, as indicated by the NASA-TLX questionnaire. Moreover, learning curves were highly heterogeneous across individuals, suggesting variability in the learning process, possibly because some participants required more repetitions or a longer training period than others. This raises questions about the feasibility of relying on this signal for everyday motor augmentation applications, especially considering the time and effort required for training.

## Limitations

While the physiological changes we observed were primarily peripheral, our ability to detect cortical adaptations was constrained by the poor sensitivity and low signal-to-noise ratio of EEG. Moreover, participants showed highly heterogeneous learning curves, with some achieving higher scores in the asynchronous control phase than others, alongside corresponding differences in beta band activity changes. Some participants did not improve their control at all. Although a group level trend was present, the response to the 10-day training protocol was clearly highly variable across individuals. This heterogeneity may reflect differences in the number of repetitions needed for learning, or the demanding nature of the protocol, which was lengthy and required sustained attention.

## Conclusion

We have shown that participants can learn to voluntarily increase peripheral beta band power through longitudinal neurofeedback training. This training induced physiological adaptations at the peripheral level, including increased beta band power and strengthened common synaptic input to the motor neuron pool, as reflected by intramuscular coherence. These adaptations were specific to the beta band, with no comparable changes observed in the delta, theta, or alpha bands. Furthermore, the observed changes were confined to the periphery, as neither cortical beta band activity nor corticomuscular coherence changed following training. Together, these findings demonstrate that peripheral beta band activity is a dynamic neural oscillation that can be shaped through targeted behavioural training.

These findings are relevant in several contexts. Abnormal beta band power and corticomuscular coupling are observed across multiple neurological conditions, yet the mechanisms underlying the transition from healthy to pathological states remain unclear. Our results suggest that these changes may arise from other neural mechanisms rather than from a cortical source. In addition, because peripheral beta band activity can be decoded non-invasively, independently of force generation, and progressively brought under voluntary control, it may represent a promising candidate for future neural interfaces and motor augmentation applications.

